# Lactoferrin as potential supplementary nutraceutical agent in COVID-19 patients: *in vitro* and *in vivo* preliminary evidences

**DOI:** 10.1101/2020.08.11.244996

**Authors:** Elena Campione, Caterina Lanna, Terenzio Cosio, Luigi Rosa, Maria Pia Conte, Federico Iacovelli, Alice Romeo, Mattia Falconi, Claudia Del Vecchio, Elisa Franchin, Maria Stella Lia, Marilena Minieri, Carlo Chiaramonte, Marco Ciotti, Marzia Nuccetelli, Alessandro Terrinoni, Andrea Magrini, Sergio Bernardini, Massimo Andreoni, Loredana Sarmati, Alessandro Miani, Prisco Piscitelli, Piera Valenti, Luca Bianchi

**Affiliations:** Dermatology Unit, Tor Vergata University Hospital, Department of Systems Medicine, Rome, 00133, Italy; Department of Public Health and Infectious Diseases, University of Rome “La Sapienza”, 00185, Italy; Department of Biology, Structural Bioinformatics Group, University of Rome “Tor Vergata”, Rome, 00133, Italy; Department of Molecular Medicine, University of Padova, 35122 Padova, Italy; Department of Experimental Medicine, Tor Vergata University Hospital, Rome, 00133, Italy; Department of statistics, University of Rome Tor Vergata, Rome, 00133, Italy; Virology Unit, Tor Vergata University Hospital, Rome, 00133, Italy; Laboratory Medicine, Department of Experimental Medicine and Surgery, Tor Vergata University Hospital; Occupational Medicine Department, University of Rome “Tor Vergata”, Rome, 00133, Italy; Villa dei Pini Hospital, Anzio (RM), Italy; Pineta Grande Hospital, Caserta, Italy; Fimmg provincial, Rome, Italy; Infectious Disease Unit, Tor Vergata University Hospital, Rome, 00133, Italy; Department of Environmental Sciences and Policy, University of Milan, Via Celoria 2, 20133 Milan, Italy; UNESCO Chair on Health Education and Sustainable Development, University of Naples Federico II, 80131 Naples, Italy

**Keywords:** lactoferrin, COVID-19, SARS-CoV2

## Abstract

Lactoferrin, a multifunctional cationic glycoprotein, secreted by exocrine glands and neutrophils, possesses an antiviral activity extendable to SARS-CoV-2.

We performed *in vitro* assays proving lactoferrin antiviral activity through direct attachment to both virus and cell surface components. This activity varied according to concentration (100/500μg/ml), multiplicity of infection (0.1/0.01) and cell type (Vero E6/Caco-2 cells).

Interestingly, the *in silico* results strongly supported the hypothesis of a direct recognition between the lactoferrin and the Spike S glycoprotein, thus hindering the viral entry into the cells.

Hence, we conducted a clinical trial to investigate effect and tolerability of a liposomal lactoferrin formulation as a supplementary nutraceutical agent in mild-to-moderate and asymptomatic COVID-19 patients.

A total of 92 mild-to-moderate (67/92) and asymptomatic (25/92) COVID-19 patients were recruited and divided in 3 groups according to the administered regimen. Thirty-two patients, 14 hospitalised and 18 in home-based insolation received oral and intranasal liposomal bovine lactoferrin (bLf), 32 hospitalised patients were treated with standard of care treatment (hydroxychloroquine, azitromicin and lopinavir/darunavir), and 28, in home-based isolation, did not take any medication. Furthermore, 32 COVID-19 negative, not-treated, healthy subjects were added as a control group for ancillary analysis.

bLf-supplemented COVID-19 patients obtained an earlier and significant (p < 0,0001.) median rRT-PCR SARS-COV-2 RNA negative conversion than standard of care-treated and non-treated COVID-19 patients (14.25 vs 27.13 vs 32.61 days, respectively).

In addition, bLf-supplemented COVID-19 patients showed significant fast clinical symptoms recovery than standard of care-treated and non-treated COVID-19 patients. Moreover, in bLf-supplemented patients, a significant decrease of either serum ferritin or IL-6 levels or host iron overload, all parameters characterizing inflammatory processes, were observed. Serum D-dimers was also found significantly decreased following bLf supplement. No adverse events were reported.

These *in vitro* and *in vivo* observations led us to speculate a potential and safe supplementary role of Blf in the management of mild-to-moderate and asymptomatic COVID-19 patients.

## INTRODUCTION

In December 2019, in Wuhan, China, a cluster of pneumonia cases was observed. This cluster was related to a novel member of *Betacoronavirus*, named SARS-CoV-2, possessing more than 80% identity to SARS-CoV and 50% to the MERS-CoV [1,2]. Coronavirus are spherical, enveloped viruses possessing a single-strand, positive-sense RNA genome ranging from 26 to 32 kilobases in length[3]. Their genome encodes 16 non-structural proteins [4], accessory proteins [5] and 4 essential structural proteins, namely spike S glycoprotein, small envelope protein, matrix protein, and nucleocapsid protein[6]. Homotrimeric S glycoprotein, possessing N-linked glycans, is located on the envelope and comprises two subunits (S1 and S2) in each spike monomer[7]. As homotrimers of S glycoproteins are exposed on the viral surface, they are responsible for binding to host receptors (S1) and membrane fusion (S2)[1,8]. Cryo-electron microscopy on S protein has highlighted its interaction with cell receptor angiotensin-converting enzyme 2 (ACE2) and the dissociation of S1 after binding to the host cells. This leads S2 to a more stable state, pivotal for membrane fusion [9–11]. Apart from ACE2, also the heparan sulfate proteoglycans (HSPGs), located on the cell surface, have been recognized as the binding sites for SARS-CoV [12] and could be important also for SARS-CoV-2 in the early attachment phase.

Lately, Wrapp and coworkers [13], determined the first 3.5 Å resolution cryo-electron microscopy (cryo-EM) structure of the SARS-CoV-2 S trimer in the prefusion conformation. Because of the critical function of S glycoprotein in the SARS-CoV-2 infection process, the knowledge of this structure, which represents a target for antibody, protein and drug mediated neutralization, allowed to get atomic-level information able to guide the design and development of innovative therapeutic molecules [14].

Considering the hypothesis that innate immunity could suggest possible molecules with antiviral activity against SARS-CoV-2, we highlighted how children, in which innate immunity is more prominent [15], are less likely to suffer of severe or critical COVID19 disease than adults [16,17]. Preliminary evidences indicated that the breastmilk from mothers with COVID-19 is free from SARS-CoV-2[15]

Infact LF could interact with both maternal and fetal microenvironments being physical as well as immunological barriers against pathogens.[16]

Lactoferrin (Lf), a multifunctional glycoprotein, belonging to the transferrin family, secreted by exocrine glands and neutrophils and present in human milk and in all secretions [17,18], could behave as potential nutraceutical agent able to counteract SARS-CoV-2 infection.

Indeed, two promising *in vitro* studies, the first on SARS-CoV [12] and the second on SARS-CoV-2 [19] have demonstrated that Lf is able to inhibit the early phase of virus infection [12] and is efficient against SARS-CoV-2 also in post-infection phase[19].

The pleiotropic activity of Lf is mainly based on its four different functions: to chelate two ferric irons per molecule, to interact with anionic molecules, to enter inside the nucleus and to modulate iron and inflammatory homeostasis. The ability to chelate two ferric ions per molecule is associated to the inhibition of reactive oxygen species formation and the sequestration of iron, which is important for bacterial and viral replication and is at the basis of the antibacterial and antiviral activity of Lf [18,20,21]. The binding of Lf to the anionic surface compounds, thanks to Lf cationic feature, is associated to the host protection against bacterial and viral adhesion and entry [18]. Moreover, the entrance of Lf inside host cells and its translocation into the nucleus [22,23] is related to the anti-inflammatory activity of Lf [24–26]. Furthermore, its ability to restore iron homeostasis, perturbed by viral infection and inflammation [27], is associated to its ability to chelate iron, to decrease iron overload, to diminish IL-6 levels and to modulate iron proteins. As a matter of fact, iron homeostasis involves several iron proteins such as transferrin, ferroportin, hepcidin and ferritin and their disorders of which, induced by inflammation, lead to intracellular iron overload which favours viral replication [28]. Moreover, Lf seems to regulate the activation of plasminogen and to control the coagulation cascade with a remarkable antithrombotic activity[29], a very frequent complication of SARS-CoV2 infection [30]. In addition to all these abilities, Lf, as above reported, inhibits the early phase of SARS-CoV[12] and the post-infection phase of SARS-CoV-2 [12,19].

Therefore, based on these informations, and in order to evaluate the possibility of using Lf in the clinical treatment of COVID-19, we tested the antiviral activity in *in vitro* experiments to verify if its activity was related to the binding to viral particles and/or to host cells similarly to what happens with other viruses [22,23]. Furthermore, the SARS-CoV-2 S trimer structure in prefusion conformation [13] has been used to perform a protein-protein molecular docking analysis with the aim to confirm the hypothesis of a direct interaction between the S glycoprotein and the Lf protein. The structure of the Spike glycoprotein [13] has been completed using modelling techniques and used to predict Lf interaction sites. Furthermore, the selected high-score protein-protein complex has been structurally investigated using classical molecular dynamics (MD) simulation and the free energy of interaction between these proteins has been evaluated through the molecular mechanic energies combined with generalized Born and surface area continuum solvation (MM/GBSA) method[31].

Hence, a clinical trial has been designed to validate the aforementioned assumptions. We conducted a parallel 3 groups clinical trial to investigate effect and tolerability of a liposomal bLf formulation as a supplementary nutraceutical agent in COVID19 patients. A total of 92 COVID19 patients, 25/92 asymptomatic and 67/92 mild-to-moderate, were recruited and divided in 3 groups according to the administered regimen: 32/92 COVID-19 patients, 14 hospitalised and 18 in home-based isolation, received oral and intranasal liposomal bLF supplement; 32 COVID-19 hospitalised patients were treated with hydroxychloroquine, azitromicin and lopinavir/darunavir as standard of care treatment (SOC); twenty eight COVID-19 patients, in home-based isolation did not take any medication against COVID-19. Furthermore, a group of 32 healthy subjects with negative COVID19 rRT-PCR was added as a control group for ancillary analysis.

## MATERIALS & METHODS

### In vitro antiviral activity of lactoferrin

For *in vitro* experiments, highly purified bovine Lf (bLf) was kindly provided by Armor Proteines Industries (France). BLf was checked by SDS-PAGE and silver nitrate staining. Its purity was about 98% and its concentration was confirmed by UV spectroscopy according to an extinction coefficient of 15.1 (280 nm, 1% solution). The bLf iron saturation was about 7% as detected by optical spectroscopy at 468 nm based on an extinction coefficient of 0.54 (100% iron saturation, 1% solution). LPS contamination of bLf, estimated by Limulus Amebocyte assay (Pyrochrome kit, PBI International, Italy), was equal to 0.6 ± 0.05 ng/mg of bLf. Before each *in vitro* assay, bLf solution was sterilized by filtration using 0.2 μm Millex HV at low protein retention (Millipore Corp., Bedford, MA, USA).

### Cell culture and virus

The African green monkey kidney-derived Vero E6 and human colon carcinoma-derived Caco-2 cells were provided by American Type Culture Collection (ATCC). Cells were grown in high-glucose Dulbecco’s modified Eagle’s medium (DMEM) (Euroclone, Milan, Italy) supplemented with 10% fetal bovine serum (FBS) (Euroclone, Milan, Italy) at 37°C in humidified incubators with 5% CO_2_. SARS-CoV-2 strain was isolated from nasopharyngeal specimen taken from a patient with laboratory confirmed COVID-19 and was propagated in Vero E6 cells. Viral titres were determined by 50% tissue culture infectious dose (TCID50) assays in Vero E6 (Spearman-Kärber method) by microscopic scoring. All experiments were performed by infecting Vero E6 and Caco-2 cells with SARS-CoV-2 strain at the Department of Molecular Medicine, University of Padua, under Biosafety Level 3 (BSL3) protocols, in compliance with laboratory containment procedures approved by the University of Padua.

### Cytotoxicity assay

Cytotoxicity was evaluated by incubating 100 and 500 μg of bLf - the concentrations used for *in vitro* experiments - in DMEM containing 10% of FBS for 72 h at 37°C with Vero E6 and Caco-2 cells in 96-well microtiter plates. Cell proliferation and viability were assessed by MTT assay (Merck, Italy). Tetrazolium salts used for quantifying viable cells were cleaved to form a formazan dye, which was evaluated by spectrophotometric absorbance at 600 nm.

### Infection assay

For infection assay, Vero E6 cells were seeded in 24-well tissue culture plates at a concentration of 1×10^5^ cells/well for 24h at 37°C in humidified incubators with 5% CO_2_, while Caco-2 cells were seeded at a concentration of 2×10^5^ cells/well for 48h at 37°C in humidified incubators with 5% CO_2_. In order to assess the putative inhibition of SARS-CoV-2 strain infection on Vero E6 monkey cells, 100 μg/ml of bLf were used. Conversely, the supposed antiviral activity against SARS-CoV-2 strain on Caco-2 human cells has been investigated using not only 100 but also 500 μg/ml of bLf. In order to investigate the putative interaction of bLf with viral particles and/or host cells, the following different experimental approaches in both Vero E6 and Caco-2 cells were performed. To evaluate if bLf can interfere with the viral infectivity rate by binding viral surface components, SARS-CoV-2 at multiplicity of infection (MOI) of 0.1 and 0.01 was pre-incubated with bLf for 1h at 37°C in humidified incubators with 5% CO_2_. The cells were then infected with these suspensions for 1h at 37°C in humidified incubators with 5% CO_2_. In order to evaluate if bLf interferes with the viral attachment to host cells, the cells were pre-incubated in culture medium without FBS with bLf for 1h at 37°C in humidified incubators with 5% CO_2_. The cells were then washed with phosphate buffered saline (PBS) and infected with SARS-CoV-2 at MOI of 0.1 and 0.01 for 1h at 37°C in humidified incubators with 5% CO_2_. To assess if bLf can interfere with both viral and host cell components, bLf was added together with SARS-CoV-2 at MOI of 0.1 and 0.01 to cell monolayer for 1h at 37°C in humidified incubators with 5% CO_2_. In addition, the pre-incubation of SARS-CoV-2 with bLf for 1h at 37°C was used to infect cell monolayer previously pre-treated with bLf for 1 h at 37°C.

Regarding Vero E6 cells, after each experimental approach, the cells were washed with PBS, overlaid with DMEM containing 0.75% of carboxymethylcellulose and 2% of FBS and incubated for 48h at 37°C in humidified incubators with 5% CO_2_. After 48h, the cells were washed, fixed with 5% of formaldehyde for 10 min at room temperature and stained with crystal violet at 1% for 5 min. The number of plaques was determined after extensive washing.

The other infection experiments were carried out with Caco-2 cells. Substantial cell death was not detected up to 7 days on Caco-2 cells after SARS-CoV-2 infection at MOI 0.1[32]. In this respect, after each experimental procedure, the cell monolayers were replaced with DMEM with 2% of FBS and after 6, 24 and 48 h post-infection (hpi) the supernatant samples were collected for RNA extraction and quantitative real-time reverse transcription (rRT)-PCR analysis of viral particles. Briefly, we lysed 200 μl of supernatant in an equal volume of NUCLISENS easyMAG lysis buffer (Biomerieux, France). Detection of SARS-CoV-2 RNA was performed by an in-house real-time RT–PCR method, which was developed according the protocol and the primers and probes designed by Corman et al. [33] that targeted the genes encoding envelope (E) (E_Sarbeco_F, E_Sarbeco_R and E_Sarbeco_P1) of SARS-CoV-2. Quantitative rRT–PCR assays were performed in a final volume of 25 μl, containing 5 μl of purified nucleic acids, using One Step Real Time kit (Thermo Fisher Scientific) and run on ABI 7900HT Fast Sequence Detection Systems (Thermo Fisher Scientific). Cycle threshold (Ct) data from rRT–PCR assays were collected for E genes. Genome equivalent copies per ml were inferred according to linear regression performed on calibration standard curves.

### Protein-protein docking methods

The structure of the SARS-CoV-2 Spike glycoprotein in prefusion conformation was extracted from a clustering procedure carried out in a previously published paper[14]. The three-dimensional structure of the diferric forms of bLf and hLf, refined at 2.8 and 2.2 Å resolution respectively, were downloaded from the PDB Database (PDB IDs: 1BLF[34], and 1B0L,[35]). The protein-protein docking analysis between the modelled SARS-CoV-2 Spike glycoprotein[14] and the Lf structures was carried out using the Frodock docking algorithm [36]. Frodock’s approach combines the projection of the interaction terms into 3D grid-based potentials and the binding energy upon complex formation, which is approximated as a correlation function composed of van der Waals, electrostatics and desolvation potential terms. The interaction-energy minima are identified through a fast and exhaustive rotational docking search combined with a simple translational scanning [37]. Both docking procedures were performed using Frodock’s (http://frodock.chaconlab.org/) web-server.

#### Molecular dynamics

Topology and coordinate files of the input structures were generated using the tLeap module of the AmberTools 19 package [39]. The Spike glycoprotein and Lf were parametrized using the ff19SB force field, and were inserted into a rectangular box of TIP3P water molecules, with a minimum distance of 12.0 Å from the box sides, and after neutralizing the solution with 0.15 mol/L of NaCl ions. To remove unfavourable interactions, all structures underwent four minimization cycles, each composed by 500 steps of steepest descent minimization followed by 1500 steps of conjugated gradient minimization. An initial restraint of 20.0 kcal • mol^−1^ • Å^−2^ was imposed on protein atoms and subsequently reduced and removed in the last minimization cycle. Systems were gradually heated from 0 to 300 K in a NVT ensemble over a period of 2.0 ns using the Langevin thermostat, imposing a starting restraint of 0.5 kcal • mol^−1^ • Å^−2^ on each atom, which was decreased every 500 ps in order to slowly relax the system. The systems were simulated in an isobaric-isothermal (NPT) ensemble for 2.0 ns, imposing a pressure of 1.0 atm using the Langevin barostat and fixing the temperature at 300 K. Covalent bonds involving hydrogen atoms were constrained using the SHAKE algorithm [40]. A production run of 30 ns was performed for with a timestep of 2.0 fs, using the NAMD 2.13 MD package [41]. The PME method was used to calculate long-range interactions, while a cut-off of 9.0 Å was set for short-range interactions. System coordinates were saved every 1000 steps.

#### Trajectory analysis

Distance analysis was performed using the distance module of the GROMACS 2019 analysis tools [42], while hydrogen bond persistence was evaluated using the hbonds module coupled to in-house written codes. The hydrophobic contacts were identified using the contact_map and contact_frequency routines of the mdtraj Python library [43]. Generalized Born and surface area continuum solvation (MM/GBSA) analysis were performed over the last 15 ns of the trajectories, using the MMPBSA.py.MPI program implemented in the AMBER16 software [44] on 2 nodes of the ENEA HPC cluster CRESCO6 [45]. Pictures of the Spike-Lf and Spike RBD-ACE2 complexes were generated using the UCSF Chimera program [46].

## Clinical trial

### Study population

A clinical trial has been conducted to investigate effect and tolerability of an oral and intranasal liposomal bLf formulation as supplementary nutraceutical agent in mild-to-moderate and asymptomatic COVID-19 patients. From April 2020 to June 2020 a total of 92 patients with confirmed COVID19 infection at rRT-PCR nasopharyngeal swab were recruited to participate in the study protocol. Patients were divided in three groups: 32 patients, 14 hospitalised and 18 in home-based isolation, received oral and intranasal liposomal bLF, 32 hospitalised patients were treated with SOC regimen (hydroxychloroquine, azitromicin and lopinavir/darunavir) only, 28, in home-based isolation, did not receive any anti-COVID-19 treatment.

All the recruited patients presented mild-to-moderate symptoms or were asymptomatic. A mild-to-moderate disease is defined by less severe clinical symptoms with no evidence of pneumonia and not requiring Intensive Care Unit (ICU) [38] A control group of 32 healthy volunteers, with negative rRT-PCR at the naso-oropharingeal swab, was included in the study for ancillary analysis.

### Enrollment criteria

Eligible patients were over 20 years old, with a confirmed COVID-19 rRT-PCR at the naso-oropharingeal swab and blood oxygen saturation (SPO2) > 93% or Horowitz index (PaO2 / FiO2) > 300mmHg. Patients had not previously been treated against SARS-CoV-2. Exclusion criteria included pregnancy and breastfeeding, nitric oxide and nitrates assumptions, known allergy to milk proteins, a medical history of bronchial hyperactivity or pre-existing respiratory diseases. ICU COVID-19 in-patients were excluded.

In the clinical trial, performed in the middle phase of the pandemic, placebo and liposome arms have not been included due to the specific request of our Hospital Ethics Committee.

All patients gave written informed consent after receiving an extensive disclosure of the study purposes and risks. The trial was approved by the Tor Vergata University Hospital Ethics Committee (Code 42/20). It was registered at www.clinicalTrials.gov (NCT04475120) and reported according to CONSORT guidelines (Figure S1, supplemental data).

### Patient Groups

Thirty two patients (14 hospitalised and 18 in home-based isolation) belonging to the first group received oral and intranasal liposomal bLf. BLf capsules for oral use contained 100 mg of bLf encapsulated in liposome while bLf nasal spray had about 8 mg/ml of bLf encapsulated in liposome. BLf, contained in both products, was checked by SDS-PAGE and silver nitrate staining and its purity was about 95%. The bLf iron saturation was about 5% as detected by optical spectroscopy at 468 nm based on an extinction coefficient of 0.54 (100% iron saturation, 1% solution). The scheduled dose treatment of liposomal bLf for oral use was 1gr per day for 30 days (10 capsules per day) in addition to the same formulation intranasally administered 3 times daily (a total of about 16 mg/nostril) Thirty two hospitalized patients belonging to the second group were only treated with SOC regimen according to the national guidelines at the time of the enrollment: lopinavir/ritonavir cps 200/50 mg, 2×2/day (alternatively darunavir 800 mg 1 cp/day+ritonavir 100 mg 1 cp/day or darunavir/cobicistat 800/150 mg 1 cp/day), chloroquine 500 mg, 1×2/day or hydroxychloroquine cp 200 mg, 1×2/day. SOC regimen lasted from 5 to 20 days, with timing to be established according to clinical evolution.

Twenty-eight patients, in home-based isolation, belonging to the third group did not receive any therapy.

A control group, comprising 32 healthy volunteers, did not receive any treatment or placebo.

Blood samples and clinical assessments were evaluated at baseline (T0), after 15 days (T1) and after 30 days (T2).

### Primary endpoint

Primary endpoint was to assess the mean time length needed to reach a rRT-PCR negative conversion rate of SARS-COV-2 RNA in each group of COVID-19 patients enrolled in the study.

### Secondary endpoints

Secondary endpoint was to estimate the proportion of patients who achieved disease remission defined as symptoms recovery and improvement of those blood parameters eventually deranged in COVID-19 patients.

In addition, safety and tolerability of liposomal bLf for oral and intranasal use were assessed.

### Tertiary endpoint

Tertiary endpoint was an ancillary analysis to evaluate in bLf-supplemented COVID-19 patients iron, transferrin and ferritin variation, together with the following inflammatory cytokines and markers, IL-6, IL-10, TNF-α, adrenomedullin serum levels at T0 and T2, but also in comparison with healthy volunteers’ group.

### Statistical analysis

For *in vitro* experiments, the number of plaque forming units (pfu)/ml of SARS-CoV-2 on Vero E6 cells and the number of SARS-CoV-2 RNA copies/ml on Caco-2 cells in each experimental approach was compared with the control ones (untreated SARS-CoV-2 and cells) at the same time point in order to assess the statistically significant differences by using unpaired student’s *t* tests. Results are expressed as the mean values ± standard deviation (SD) of three independent experiments. In each case, a *p* value ≤ 0.05 was considered statistically significant.

For what concerns the clinical trial, descriptive and inferential statistical analyses were performed. The Kolmogorov–Smirnov test was used to check the normal distribution of blood parameters.

Blood parameters obtained at T0 in COVID-19 group and control group were compared using t-test. Data were then analyzed with a significant two-tailed p-value ≤ 0.05.

All parameters obtained at T0 and T2 in COVID-19 group were then compared using paired t-test. In addition, the mean change between T0 and T2 was also assessed using paired t-test. Normally distributed data were then analyzed with a significant p-value ≤ 0.05.

## RESULTS

### Lactoferrin displays antiviral properties in *in vitro* models

Preliminary, the doses of bLf in native form (7% iron saturated) corresponding to 100 μg/ml for Vero E6 cells and 100 and 500 μg/ml for Caco-2 cells were assayed to detect their putative cytotoxicity by measuring cell morphology, proliferation and viability after 72 h of incubation. Both 100 and 500 μg/ml of bLf do not exert any cytotoxic effect (data not shown).

Then, the efficacy of different concentrations of bLf in inhibiting SARS-CoV-2 infection was tested on Vero E6 and Caco-2 cells according to different experimental procedures: I) control: untreated SARS-CoV-2 and cells; II) bLf pre-incubated with virus inoculum for 1 h at 37°C before cell infection; III) bLf pre-incubated with cells for 1 h at 37°C before virus infection; IV) bLf added together with virus inoculum at the moment of infection step; V) virus and cells separately pre-incubated with bLf for 1 h at 37°C before infection.

The results obtained with Vero E6 cells are shown in Figure 1A (MOI 0.1) and 1B (MOI 0.01).

**Figure 1.**
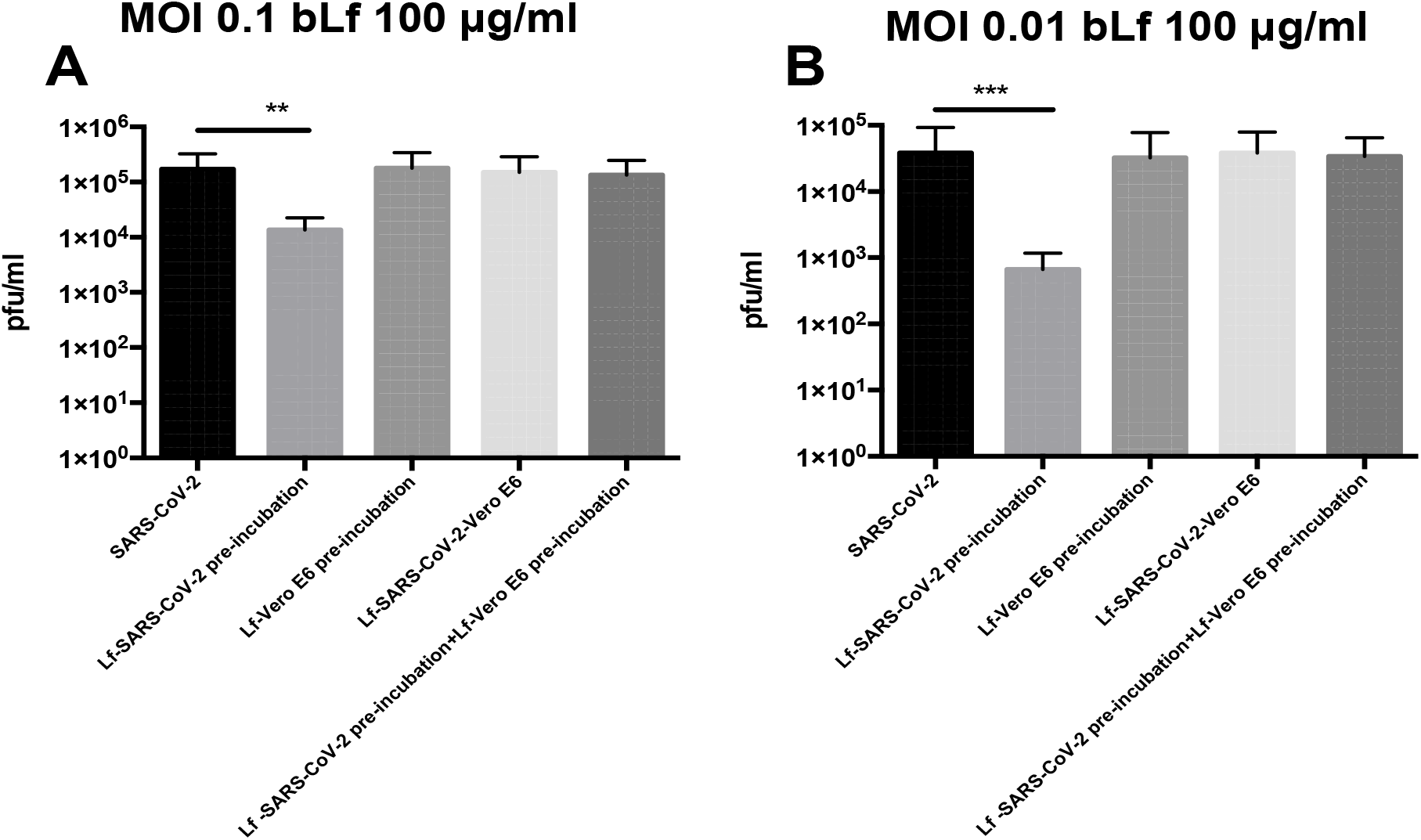
Plaque forming units (pfu)/ml of SARS-CoV-2 observed in Vero E6 cells infected at multiplicity of infection (MOI) of 0.1 (**A**) and 0.01 (**B**) in the presence or absence of 100 μg/ml of bovine lactoferrin (bLf) according to the following experimental procedures: i) control: untreated SARS-CoV-2 and Vero E6 cells; ii) bLf pre-incubated with SARS-CoV-2 inoculum for 1h at 37°C before cell infection iii) cells pre-incubated with bLf for 1 h at 37°C before SARS-CoV-2 infection; iv) bLf added together with SARS-CoV-2 inoculum during the adsorption step; v) virus and cells separately pre-incubated with bLf for 1 h at 37°C before infection. Data represent the mean values of three independent experiments. Error bars: standard error of the mean. Statistical significance is indicated as follows: **: *p* < 0.001, ***: *p* < 0.0001 (Unpaired student’s *t* test).

Regarding Vero E6 cells, an inhibition of SARS-CoV-2 replication of about 1 log at MOI 0.1 and about 2 log at MOI 0.01 on cell monolayers was observed when 100 μg/ml of bLf were pre-incubated for 1 h with virus before infection compared to untreated SARS-CoV-2 infection (*p* < 0.001 and *p* < 0.0001, respectively) (Figure 1A and 1B).

On the contrary, the data illustrated in Figure 1A and 1B, independently from the MOI used, indicate that bLf, at this concentration, does not block SARS-CoV-2 infection when it is pre-incubated with Vero E6 cells or when bLf is contemporary added to viral particles and cells at the moment of infection (Figure 1A, 1B). BLf is also ineffective when it is pre-incubated for 1 h at 37°C separately with virus and cells before infection (Figure 1A, 1B).

The efficacy of 100 and 500 μg/ml of bLf against SARS-CoV-2, assayed in Caco-2 cells, is showed in Figure 2A and 2B (MOI 0.1) and in Figure 2C and 2D (MOI 0.01), respectively.

**Figure 2:**
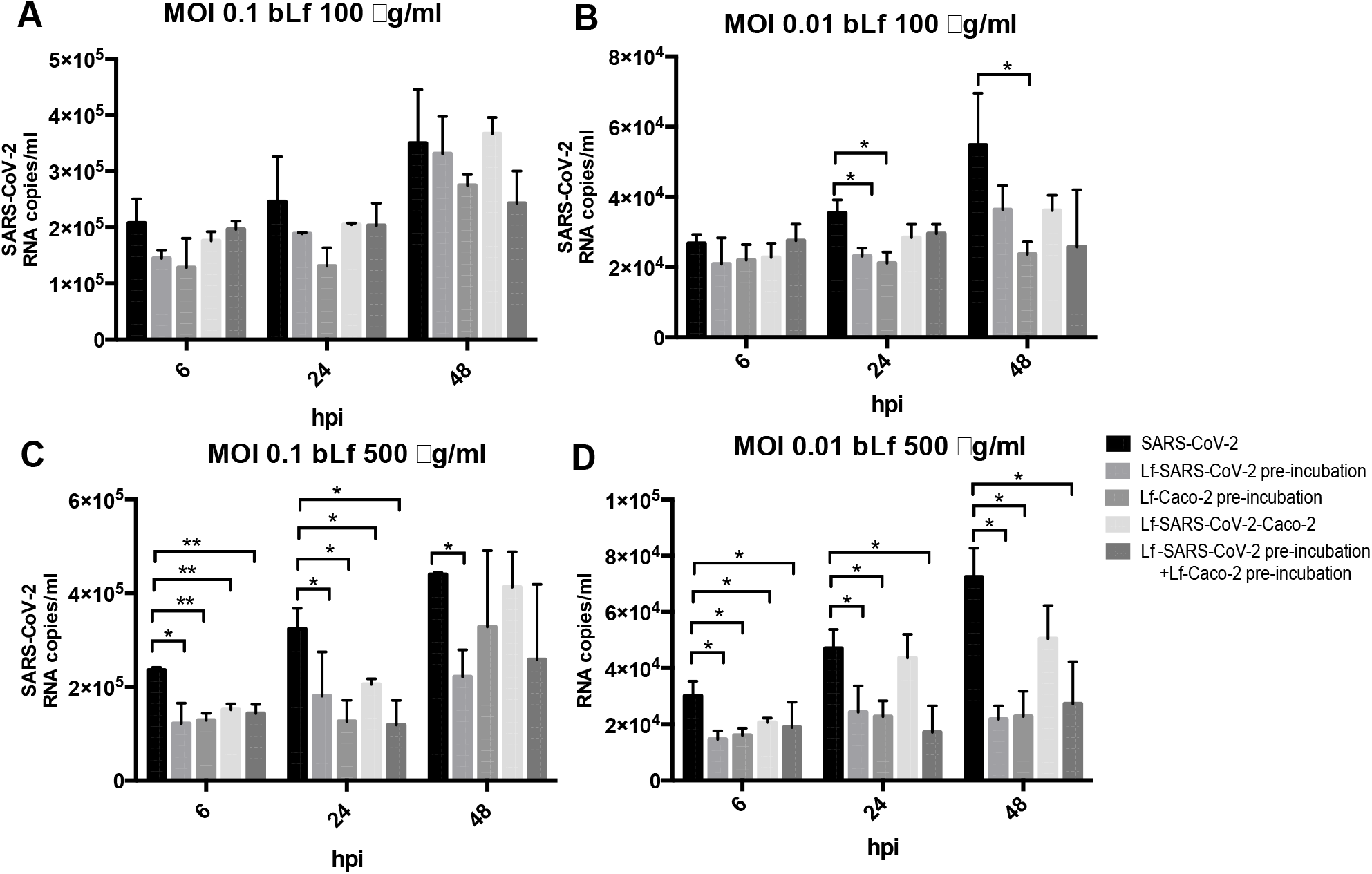
RNA copies/ml of SARS-CoV-2 observed in supernatants of Caco-2 cells infected at multiplicity of infection (MOI) of 0.1 (**A,C**) and 0.01 (**B,D**) in the presence or absence of 100 μg/ml (**A,B**) and 500 μg/ml (**C,D**) of bovine lactoferrin (bLf) according to the following experimental procedures: i) control: untreated SARS-CoV-2 and Caco-2 cells; ii) bLf pre-incubated with SARS-CoV-2 inoculum for 1h at 37°C before cell infection; iii) cells pre-incubated with bLf for 1 h at 37°C before SARS-CoV-2 infection; iv) bLf added together with SARS-CoV-2 inoculum during the adsorption step; v) virus and cells separately pre-incubated with bLf for 1 h at 37°C before infection. Viral supernatant samples were harvested at 6, 24 and 48 hours post infection (hpi). Viral loads were ascertained with quantitative rRT-PCR. Data represent the mean values of three independent experiments. Error bars: standard error of the mean. Statistical significance is indicated as follows: *: *p* < 0.05, **: *p* < 0.001 (Unpaired student’s *t* test).

Regarding Caco-2 cells, at MOI 0.1, no significant differences were observed in all experimental conditions compared to the control ones when using bLf at 100 μg/ml (Figure 2A). At MOI 0.01, an inhibition of viral load in supernatants was observed at 24 hpi only when 100 μg/ml of bLf was pre-incubated with the viral inoculum and when the cells were pre-incubated with 100 μg/ml of bLf compared to the control one (*p* < 0.05) (Figure 2B). At 48 hpi, an inhibition of viral load was observed only when the cells were pre-incubated with bLf (*p* < 0.05) (Figure 2B).

When bLf was used at a concentration of 500 μg/ml, a decrease of viral load up to 48 hpi was observed when the viral inoculum was pre-incubated with bLf compared to the control group, independently from the MOI used (*p* < 0.05) (Figure 2C, 2D). When the cells were pre-incubated with bLf, a decrease of viral load up to 24 hpi was observed compared to the control at MOI 0.1 (*p* < 0.001 after 6 hpi and *p* < 0.05 after 24hpi) (Figure 2C), while at MOI 0.01 the decrease of viral load remained statistically significant up to 48 hpi compared to the control group (*p* < 0.05) (Figure 2D). When bLf was added together with SARS-CoV-2 inoculum during the adsorption step a decrease of viral load up to 24 hpi was observed compared to untreated SARS-CoV-2 infection, independently from the MOI used (*p* < 0.001 after 6 hpi and *p* < 0.05 after 24hpi for MOI 0.1; *p* < 0.05 after 6 and 24 hpi for MOI 0.01) (Figure 2C, 2D). When the cells were pre-incubated with bLf and infected with SARS-CoV-2 previously pre-incubated with bLf, a decrease of viral load up to 24 hpi was observed for MOI 0.1 compared to untreated SARS-CoV-2 infection (*p* < 0.001 after 6 hpi and *p* < 0.05 after 24hpi for MOI 0.1) (Figure 2C), while at MOI 0.01 the decrease of viral load remains statistically significant up to 48 hpi compared to untreated SARS-CoV-2 infection (*p* < 0.05) (Figure 2D).

### Computational results

The molecular docking simulation suggests a potential interaction of the bLf structure with the Spike glycoprotein CDT1 domain in the up conformation (Figure 3A). The first three solutions obtained by Frodock clustering procedure account for more than 60% of the total generated complexes, which are almost completely superimposable to that shown in Figure 3A. Starting from the first Frodock solution, we performed a 30 ns long classical MD simulation in order to verify the stability of the complex and check for the presence of persistent interactions between the two proteins. As shown in Figure S2A (supplemental data), the distance between the centers of the mass of Spike and bLf, calculated as a function of time, oscillates around the value of 4.5 nm, indicating a constant close contact between the two molecules. MM/GBSA analysis confirmed the high affinity of the bLf for the Spike CDT1 domain (Table S1A, supplemental data), showing interaction energy of −28.02 kcal/mol. In particular, MM/GBSA results highlighted that the Van der Waals term mainly contribute to the binding energy (Table S1A, supplemental data).

**Figure 3.**
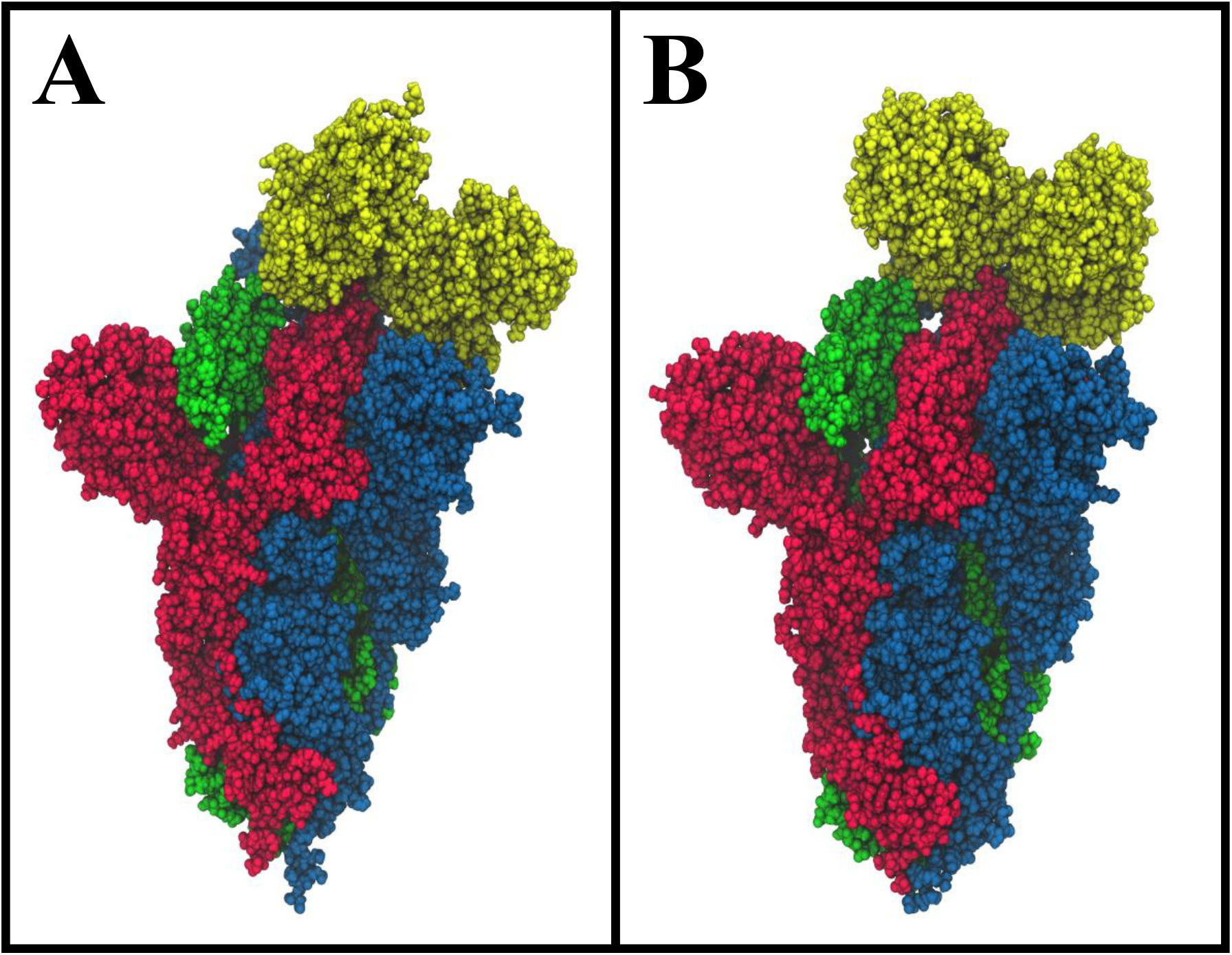
Spacefill representations of the best molecular complex obtained with Frodock between the bovine (A) and human (B) lactoferrin with the Spike glycoprotein. The red, blue and green colours represent the Spike glycoprotein chains, while the yellow depicts the lactoferrin molecules.

A detailed analysis of the interaction network reveals the presence of 28 different interactions, which persist for more than 25% of the simulation time, in agreement with the high interaction energy calculated. In detail, we found 3 salt bridges, 5 hydrogen bonds and 20 residue pairs involved in hydrophobic contacts (Table S2 left side, supplemental data).

To check if some of the Spike residues targeted by the bLf protein are involved in the binding with ACE2, we have compared the average structure extracted from the simulation with the ACE2/CDT1 domain complex structure (PDB ID: 6LZG[39] Figure 4). Surprisingly, only two Spike residues (Gly502 and Tyr505) are shared between the complexes interfaces (Table S2 left side, supplemental data), as evaluated from the inspection of the superimposed structures and from the paper analysis[39]. Despite this, Lf holds the same position assumed by the ACE2 enzyme, i.e. above the up CDT1 domain.

**Figure 4.**
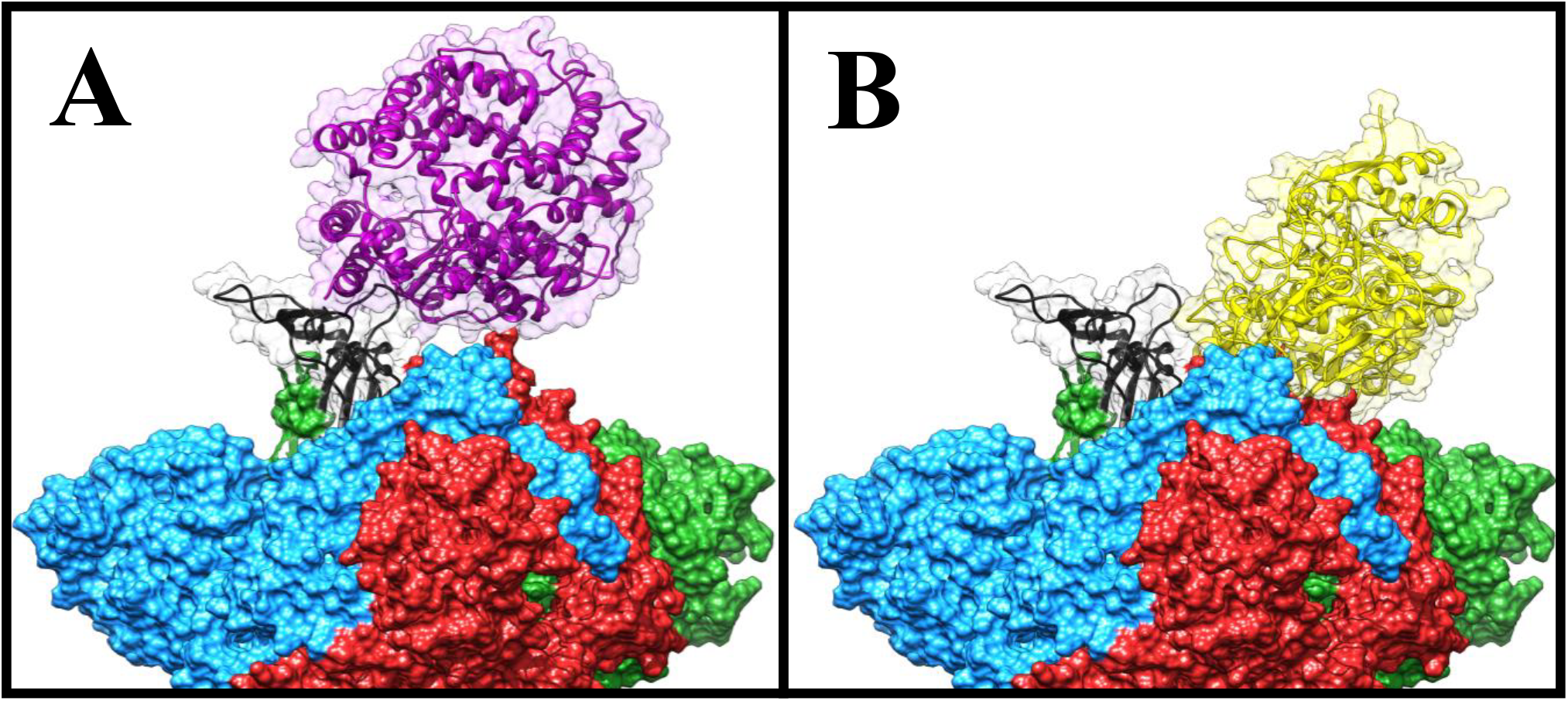
Comparison of the Frodock best complex and of the ACE2-Spike glycoprotein (PDB ID: 6LZG). The red, blue and green solid surfaces represent the three different chains composing the Spike glycoprotein. The black ribbons highlight the CTD1 domain in the up conformation. The magenta and yellow ribbons represent the ACE2 (A) and the bovine lactoferrin (B), respectively, surrounded by a transparent molecular surface representation, in order to point out the positions occupied in the space by the different structures.

We performed the same analysis over the evaluated human Lf (hLf)-Spike complex, obtaining a binding pose superimposable to that observed for the bovine protein (Figure 3B). Besides the fact that using the human protein we can still observe a persistent and close contact between the two molecules (Figure S2B, supplemental data), the analysis of the interaction network indicates the presence of a larger number of interactions (45), in agreement with a higher interaction energy revealed by the MM/GBSA approach (−48.25 kcal/mol, Table S2 right side, supplemental data). In detail, we found 12 salt bridges, 10 hydrogen bonds and 23 residue pairs involved in hydrophobic contacts (Table S1B, supplemental data), in agreement with the presence of a negative electrostatic contribution term (Table S1B, supplemental data). Comparing the average structure extracted from the simulation with the ACE2/CDT1 domain complex structure (PDB ID: 6LZG [47]) (Figure S3, supplemental data), we observed that also for the hLf only two residues (Thr500 and Tyr505) are shared between the complexes interfaces (Table S2 right side, supplemental data).

These results allow us to hypothesize that, in addition to the HSPGs binding[12], both bLf and hLf should be able to hinder the Spike glycoprotein attachment to the ACE2 receptor, consequently blocking the virus from entering into the cells.

## CLINICAL TRIAL

### Demographic data

A total of 92 patients with confirmed COVID-19 infection at rRT-PCR were recruited to participate in the study protocol (Figure 5).

**Figure 5.**
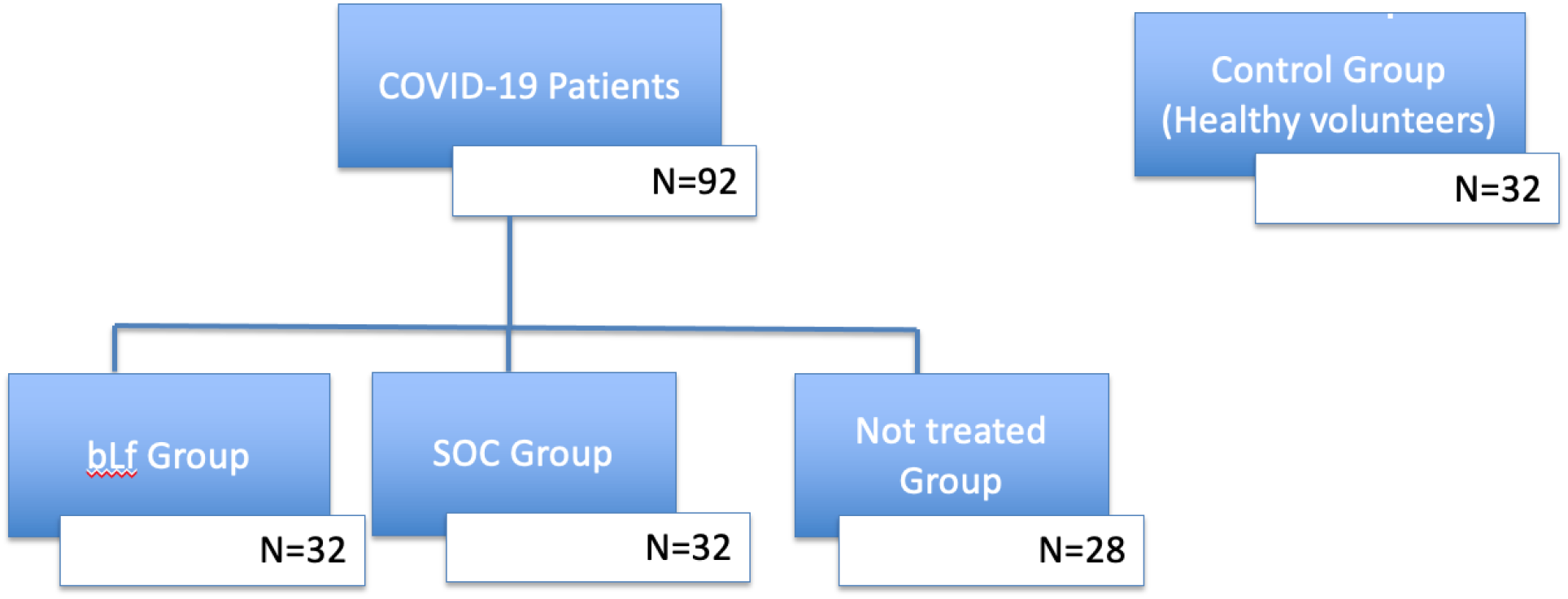
Study protocol flowchart. A total of 92 patients with confirmed COVID-19 infection at rRT-PCR were recruited to participate in the study protocol. Patients were divided in three groups according to the administered treatment. Thirty-two patients were supplemented with bLf, other 32 patients received SOC regimen, and finally 28 patients did not receive any treatment against COVID-19. A control group of 32 healthy volunteers with negative SARS-CoV2 rRT-PCR were also recruited.

Among the 32 bLf-supplemented patients, 22 had mild-to-moderate symptoms and 10 were asymptomatic. Demographic data of the groups are summarized in Table 1. The mean age was 54.6 ± 16.9 years old. Fourteen patients were males and 18 females. The most prevalent comorbidity was hypertension (28%) followed by cardiovascular diseases (15.6%) and dementia (12.5%).

**Table 1:** Demographic data.

| Demographic Data |  | bLf-supplemented COVID-19 group |  | SOC-treated COVID-19 group |  | Not treated COVID-19 group |  | CONTROL group (COVID-19 negative) |  |
| --- | --- | --- | --- | --- | --- | --- | --- | --- | --- |
|  |  | Mean<br>+/-<br>SD | N (%) | Mean<br>+/-<br>SD | N (%) | Mean<br>+/-<br>SD | N (%) | Mean<br>+/-<br>SD | N (%) |
| Age |  | 54.56<br>+/-<br>16.86 |  | 49.9<br>+/-<br>13.20 |  | 41.32<br>+/-<br>11.77 |  | 52.80<br>+/-<br>15.54 |  |
| Sex | male |  | 14 (44%) |  | 17 (53%) |  | 10 (36%) |  | 13 (41%) |
|  | female |  | 18 (56%) |  | 15 (47%) |  | 18 (64%) |  | 19 (59%) |
| Asymptomatic patients |  |  | 10 (31%) |  | 3 (10%) |  | 12 (43%) |  |  |
| Mild-to moderate patients |  |  | 22 (68.7%) |  | 29 (90%) |  | 16 (57%) |  |  |

In the group of 32 SOC-treated COVID-19, 29 had mild-to-moderate symptoms and 3 were asymptomatic. The mean age was 49.9 ± 13.2, 17 patients were males and 15 females (Table 1). The most prevalent comorbidity was hypertension (15,6%), followed by asthma (12,5%) and hypotiroidism (6,5%).

Among the 28 COVID-19 patients, in home-based isolation, not taking any drug against COVID-19, 16 had mild-to-moderate symptoms and 12 were asymptomatic. Mean age was 41.3±11.8 years old. Ten patients were male and 18 female (Table1). Furthermore, 4 out of these patients worsened their clinical profile with consequent hospitalization.

Among the 32 healthy volunteers (mean age 52.8 ±15.5 years old.) with negative rRT-PCR for SARS-CoV-2 RNA, 13 were male and 19 female (Table 1).

### Primary Endpoint Results

bLf-supplemented COVID-19 patients revealed a mean time length of rRT-PCR SARS-COV-2 RNA negative conversion of the naso-oropharingeal swab in 14.25 ± 6.0 days, shorter than that observed in SOC-treated or in non-treated COVID-19 groups (p < 0,0001).

SOC-treated COVID-19 patients showed a mean rRT-PCR SARS-COV-2 RNA negative conversion in 27.13 ± 14.4 days, whereas non-treated COVID-19 patients revealed a mean time length of negative conversion in 32.61 ± 12.2 days (Figure 6).

**Figure 6.**
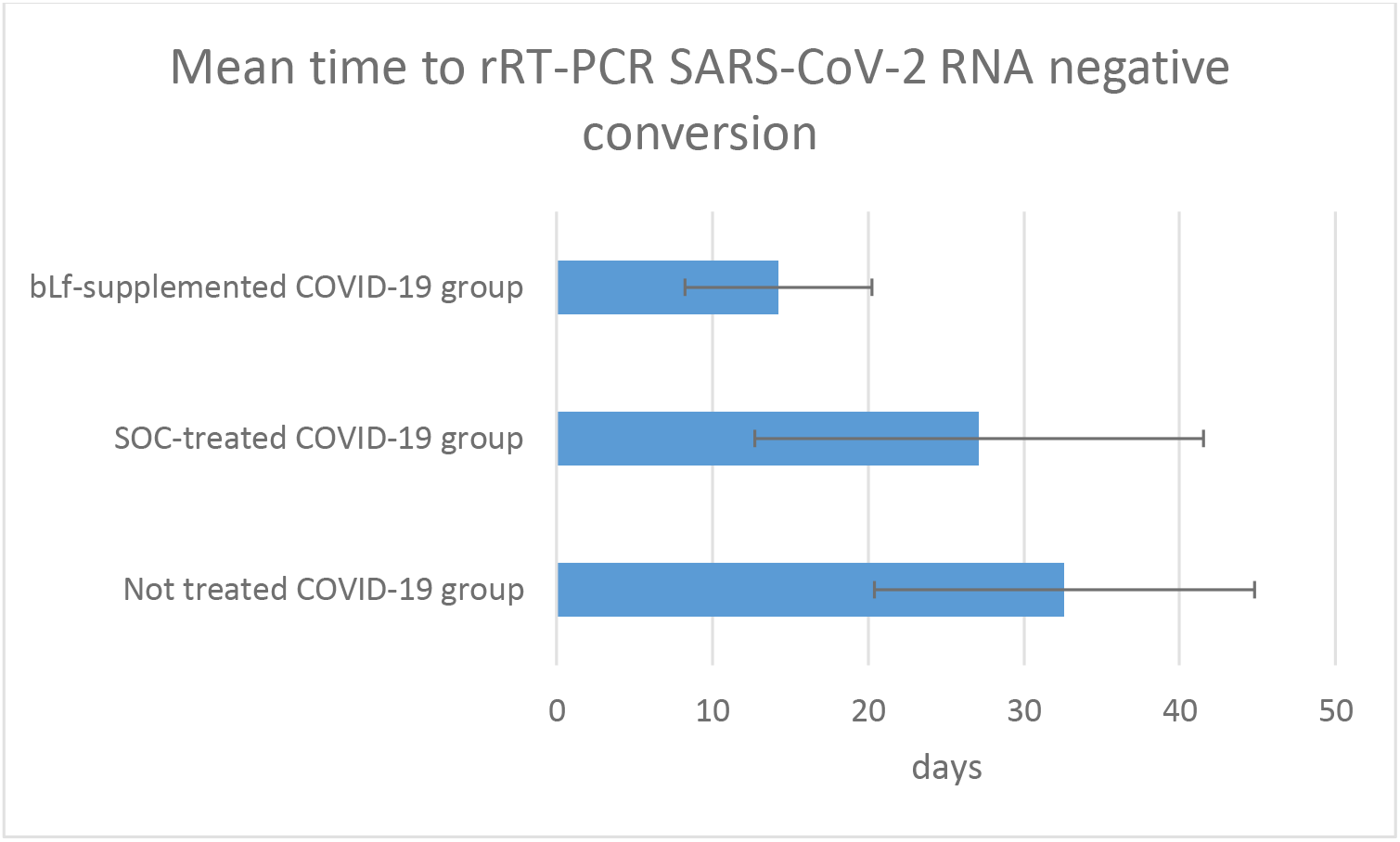
Mean time to rRT-PCR SARS-CoV-2 RNA negative conversion. bLf-supplemented COVID-19 patients revealed a mean time length of rRT-PCR SARS-COV-2 RNA negative conversion of the naso-oropharingeal swab in 14.25 ± 6.0 days; SOC-treated COVID-19 patients showed a mean rRT-PCR SARS-COV-2 RNA negative conversion in 27.13 ± 14.4 days, whereas non-treated COVID-19 patients revealed a mean time length of negative conversion in 32.61 ± 12.2 days (p < 0,0001)

### Secondary Endpoints Results

In 32 bLf-supplemented COVID-19 patients we evaluated clinical symptoms before (T0) and after therapy (T2) (Figure 7). At T0 among 32 patients, 22 were mild-to-moderate and 10 asymptomatic. The most frequent symptoms were fatigue (50%), followed by arthralgia (37.5%) and cough (28%). At T0 5 patients (15.6%) presented ageusia and 5 patients (15.6%) presented anosmia. At T1, 7 patients previously symptomatic became asymptomatic, with a total 17 asymptomatic and 15 symptomatic patients. At T2, other 6 patients, previously symptomatic at T1, became asymptomatic with a total of 23 asymptomatic and 9 symptomatic patients. The most frequent remaining symptom was fatigue followed by rare manifestations related to diarrhea, myalgia and skin manifestations. All the patients presenting anosmia and ageusia showed a complete remission of those symptoms (Figure 7).

**Fig. 7.**
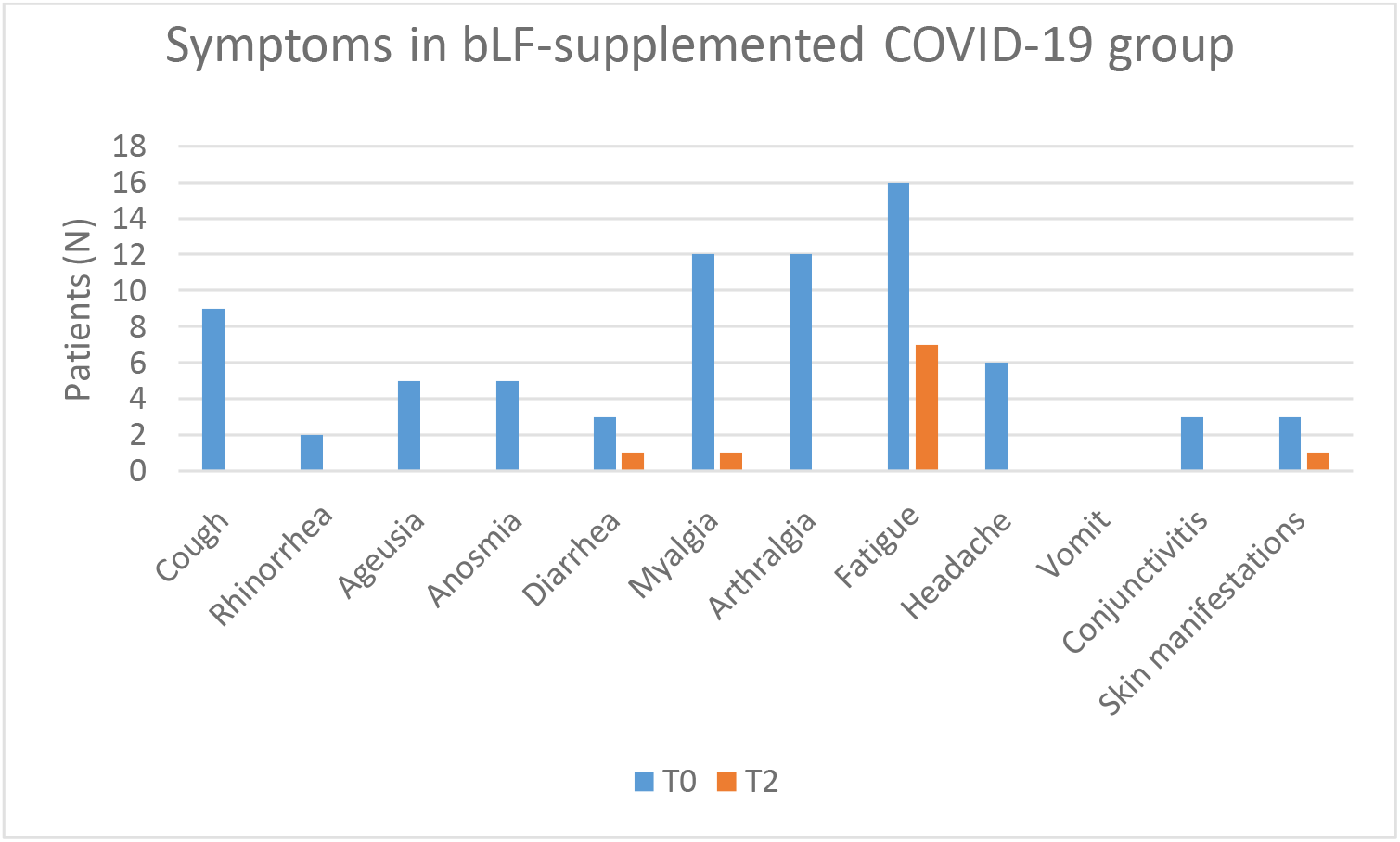
Symptoms in bLF-supplemented COVID-19 group. At T0 among the most frequent symptoms were fatigue (50%), followed by arthralgia (37.5%) and cough (28%). At T2, the most frequent remaining symptom was fatigue followed by rare manifestations related to diarrhea, myalgia and skin manifestations.

Then, we evaluated difference between blood parameters before and after bLf prescription. We noticed in bLf-supplemented group an improvement in the platelet count (T0: 239.63±83.05; T2: 243.70±65.5; ΔT2-T0 10.05±10.26) and a decrease of alanine transaminase (ALT) (T0: 29.36±22.7; T2: 23.52±12.34; ΔT2-T0 −7.32±4.36) and aspartate aminotransferase (AST) (T0: 24.36±9.80; T2: 22.64±8.33; ΔT2-T0 - 2.68±2.52). Moreover, D-dimer showed a significant decrease between T2 and T0 (ΔT2-T0 −392.56 ±142.71, p-value 0.01) in bLf-supplemented group.

Regarding safety assessment in the bLf-supplemented group, 2 patients (6.2%) showed gastrointestinal complaints related to liposomal bLf at T2. The patients did not suspend liposomal bLf and the adverse event resolved itself spontaneously.

In SOC-treated COVID-19 patients, we evaluated clinical symptoms before (T0) and after therapy (T2) (Figure 8). At T0, the most frequent symptoms were cough (72%), followed by myalgia (62%) and arthralgia (59%). Particularly, all these patients with ageusia (11) and anosmia (12) at T0 did not achieve remission at T2.

**Fig. 8.**
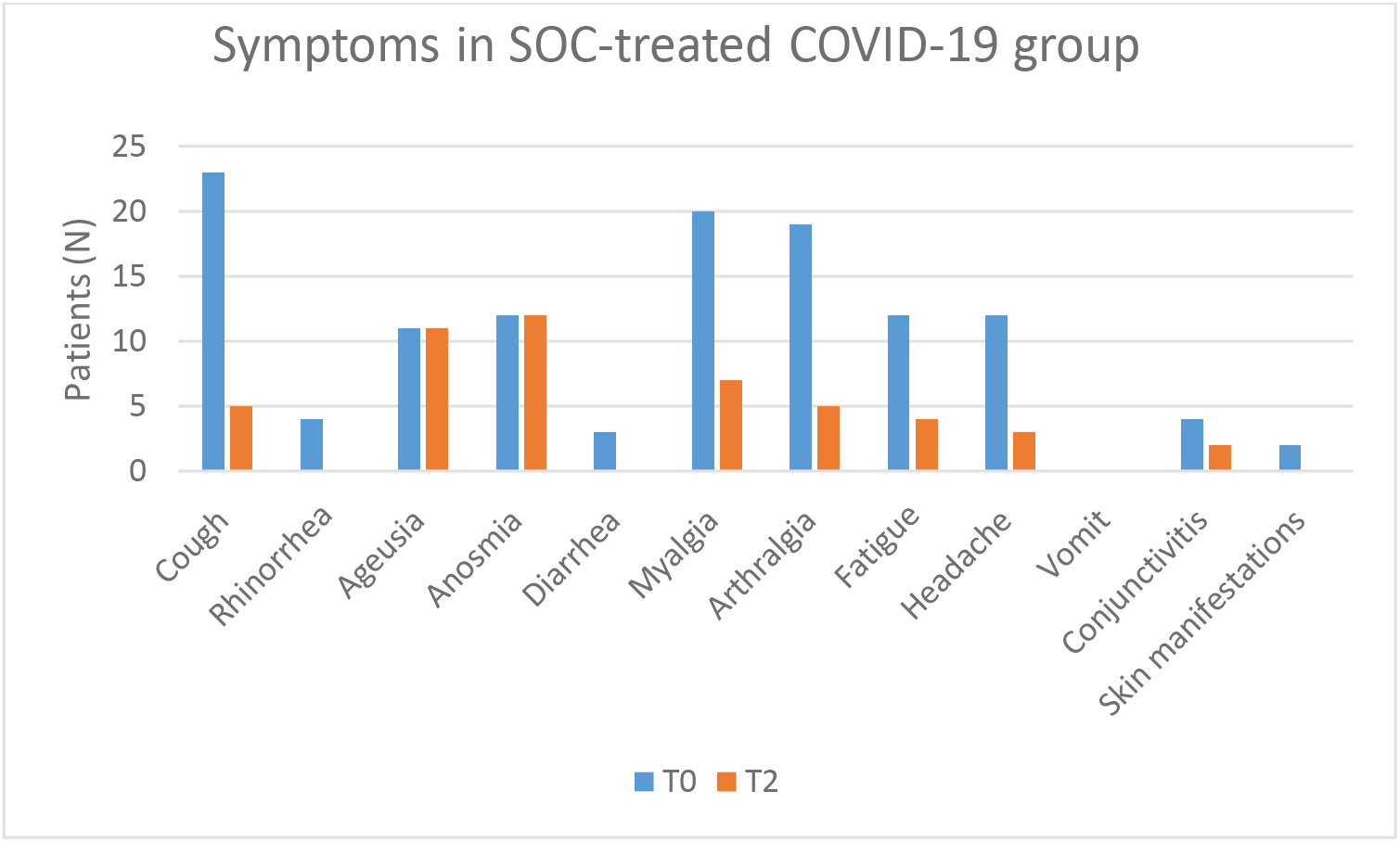
Symptoms in SOC-treated COVID-19 group. At T0, the most frequent symptoms were cough (72%), followed by myalgia (62%) and arthralgia (59%). Particularly, all these patients with ageusia (11) and anosmia (12) at T0 did not achieve remission at T2.

Then, in SOC-treated patients, we compared blood parameters before and after therapy. Among blood parameters, statistically significant variations of red blood cells (number of RBC/ml) (ΔT1 −T0 −0,43± 0,17 p-value = 0.05), hemoglobin (g/dL HGB) (ΔT1 −T0 −2,22± 0,95 p-value = 0.05), platelets count (PLT/mm^3^) (ΔT1 −T0 89,56±18,9 p-value = 0.001) were noted (Supplemental data).

#### Tertiary endpoints Results

The iron profile analysis before (T0) and after supplementation (T2) in the bLf-supplemented group revealed statistically significant values in ferritin reduction (ΔT2-T0 −90.63 ±48.49, p-value 0.04), while not statistically significance was found for serum iron and transferrin. Regarding inflammatory blood parameters, IL-6 value showed a significant decrease between T2 and T0 (ΔT2-T0 −2.52 ±1.46, p-value 0.05). Adrenomedullin remained at the same level all over the analyzed period (ΔT2-T0 − 0.01±0.03). IL-10 levels increased between T0 (8.67±3.26) and T2 (11.42±6.05), without showing statistical significance (ΔT2-T0 2.55±2.09). TNF-α decreased between T2 (25.97±21.74) and T0 (37.34 ±19.95) without showing statistical significance (ΔT2-T0 −12.92±8.81).

The comparison between bLf-supplemented COVID-19 patients and healthy volunteers group showed a significant difference in adrenomedullin (p-value< 0.0001) and IL-6 (p-value < 0.0001) (Table S3A, supplemental data).

## DISCUSSION

As still recently reported [40], there is a significant lack of treatments proven to be efficacious in early phase or mild infection, since immediate benefits of such strategies could lead to the improvement of patient outcomes and prevention of hospital recovery. Longer-term benefits may include prevention of the chronic consequences of infection as well as prevention of transmission by shortening the time length of infectiousness. Hence, in this study, we focused our attention on the well-known anti-viral and immunomodulating properties of Lf as potential supplementary nutraceutical agent in the management of mild-to-moderate and asymptomatic COVID-19 patients.

The *in vitro* antiviral activity of bLf against enveloped and naked DNA and RNA viruses has been widely demonstrated [12,20,21,41,42], while some papers have been published on its *in vivo* efficacy against viral infection [43–54].

The ability of bLf to inhibit viral infection is generally attributed to its binding to cell surface molecules and/or viral particles. BLf is able to competitively bind to heparin sulfate proteoglycans (HSPGs), present on the host cell surface and identified as initial interaction sites for enveloped viruses [55,56], thus hindering the viral adhesion and internalization [12,57,58]. Moreover, bLf can also bind directly to surface proteins of virus particles as HIV V3 loop of the gp120 [59] and HCV E2 envelope proteins [60]. The results, presented here, obtained by monitoring the effect of bLf on different experimental procedures indicate that the antiviral activity of bLf, pre-incubated with host cells, seems to vary according to MOI, different cell lines and bLf concentration. As a matter of fact, the pre-incubation of Vero E6 monolayers with 100 μg/ml of bLf, before SARS-CoV-2 infection at MOI 0.1 and 0.01, were ineffective in inhibiting virus internalization (Figure 1), differently to that observed when 100 μg/ml of bLf were pre-incubated with Caco-2 cells and the infection was performed at MOI 0.01 (Figure 2B). This antiviral activity was observed until 48 hpi.

The pre-incubation of 100 μg/ml of bLf with SARS-CoV-2 showed a significant antiviral activity higher at 0.01 MOI compared to 0.1 MOI after infection of Vero E6 cells (Figure 1A, 1B), while a significant antiviral activity assayed on Caco-2 cell lines was observed only with MOI 0.01 at 24 hpi (Figure 2B). In the other two experimental conditions, bLf did not show any significant antiviral activity on both Vero E6 and Caco-2 cells.

The pre-incubation of 500 μg/ml of bLf with Caco-2 cells showed a decrease of viral load until 24 hpi at MOI 0.1 and up to 48 hpi at MOI 0.01. Furthermore, the pre-incubation of 500 μg/ml of bLf with SARS-CoV-2 showed a significant decrease of SARS-CoV-2 RNA copies at both MOI 0.1 and 0.01. This antiviral activity persisted from 6 to 48 hpi (Figure 2C, 2D). In the other two experimental conditions, bLf exerted a significant antiviral activity only at 6 and 24 hpi when the MOI corresponded to 0.1 (Figure 2C). At MOI 0.01, a decrease of viral load up to 24 hpi was observed when bLf was added together with SARS-CoV-2 inoculum during the adsorption step (Figure 2D), while a decrease of viral load until 48 hpi was observed when both the cell monolayer and SARS-CoV-2 were previously pre-incubated with bLf (Figure 2D).

Our experimental results indicate that bLf exerts its antiviral activity either by direct attachment to the viral particles or by obscuring their cellular receptors. Moreover, the results obtained through the molecular docking and molecular dynamics simulation approaches strongly support the hypothesis of a direct recognition between the bLf and the Spike S glycoprotein. The affinity between their molecular surfaces, the large number of atomistic interactions detected and their persistence during the simulation suggest that this recognition is very likely to occur and that bLf may hinder the Spike S attachment to the human ACE2 receptor, consequently blocking the virus from entering into the cells.

Taken together these results reveal that, even if the definitive mechanism of action still has to be completely investigated, the antiviral properties of bLf are also extendable to SARS-CoV-2 virus.

In order to explore the helpful antiviral and immunomodulating effect of bLf and its possible role in the management of mild-to-moderate and asymptomatic COVID-19 patients, we designed a clinical trial to investigate the effect and safety of this nutraceutical agent.

Several studies based on COVID-19 clinical epidemiology indicate and propose the use of Lf in protecting against the virus also *in vivo*. Indeed, it has been reported that the incidence of COVID-19 in children aged 0-10 was only 0.9% in the Chinese cases, and infants developed a less severe disease form [61]. Consecutively, some authors postulated that breastfeeding or extensive use of bLf containing infant formula in this population may have protected from contagion or worst disease evolution [62,63].

We focused our research on mild-to-moderate and asymptomatic COVID-19 patients, considering them a transmission reservoir with possible evolution to the most severe disease form[64]. Li et al, analysing the viral shedding dynamics in asymptomatic and mildly symptomatic patients infected with SARS-CoV-2, observed a long-term viral shedding, also in the convalescent phase of the disease, where specific antibody production to SARS-CoV-2 may not guarantee viral clearance after hospital discharge. In their study, the median duration of viral shedding appeared to be shorter in pre-symptomatic patients (11.5 days) than in asymptomatic (28 days) and mild symptomatic cases (31 days)[65]. Accordingly, we documented a significantly reduced mean time length of rRT-PCR SARS-COV-2 RNA negative conversion in bLf-supplemented group comparing SOC-treated and non-treated patients suggesting a favouring gradual viral clearance and clinical symptoms recovery with a potential decrease in the risk of transmission and contagion.

Although there are currently rare satisfactory markers for predicting the worsening of the disease, some cytokines, including IL-6, IL-10 and TNF-α, and D-Dimer levels have been described as biomarkers related to the a severe SARS-CoV-2 infection [66–69]. In our study, we identified suitable deranged blood parameters to use as treatment target markers. Indeed, we found a statistically significant difference between bLF-supplemented COVID-19 patients and the control group of COVID-19 negative subjects, in several blood parameters, including IL-6, D-Dimer, ferritin and liver function parameters. Particularly, IL-6, D-Dimer and ferritin showed a significant decrease after bLf supplementation, possibly focusing them, if confirmed in a larger series of cases, as suitable COVID-19 treatment target markers.

Mainly, IL-6 elevation is considered to be associated with higher disease severity; IL-6 inhibitors, such as tocilizumab, have been used to treat severe COVID-19 patients[70,71]. The ability of Lf to down-regulate pro-inflammatory cytokines, such as IL-6, has already been demonstrated both in *in vitro*[72] and *in vivo* [73] models, as well as in clinical trials[74]. To our knowledge, even though in a small sample size, it should be the first evidence showing an IL-6 down-regulation in COVID-19 patients after bLf supplementation during SARS-CoV-2 infection.

We observed also a statistically significant decline in D-Dimer levels, crucial to define disease prognosis, possibly leading to a reduction in SARS-CoV-2 complications related to coagulation derangement. Recently, it has been shown that Lf can regulate the activation of plasminogen and control coagulation cascade with a remarkable antithrombotic activity [29]. This property could be relevant considering that COVID-19 is a prothrombotic disease and that the severity of the coagulation parameters impairment is related to a poor prognosis. In light of this view, SARS-CoV-2 is able to active prominent prothrombotic state rarely observed in viral diseases. Patients affected by severe COVID-19 pneumonia are at higher risk of imbalance of coagulation parameters and thus treated with low molecular weight heparin or unfractionated heparin at doses registered for prevention of venous thromboembolism [30].

Our clinical experience could lead us to speculate a potential protective and safe role of an early supplementation of bLf in COVID-19 patients in controlling the risk of a thromboembolic evolution of the disease. Lf can exert negative regulatory effects on cell migration via inhibition of plasminogen activation and through the regulation of fibrinolysis [29]. In addition, we observed an increased platelet count after bLf treatment. Indeed, COVID-19 induces thrombocytopenia as SARS-CoV-2 seems to entrap megakaryocytes and block the release of platelets. A bLf rebalanced platelet count induces COVID-19 viral clearance [75].

Ferritin, besides reflecting the status of iron stores in healthy individuals, represents also an acute-phase-protein up-regulated and elevated in both infectious and non-infectious inflammation. In COVID-19, it has been reported to be relevant for assessing disease severity and patients outcome [76,77]. In particular, serum ferritin concentration shows significant higher values in COVID-19 patients with worse outcome compared to good outcome[78].

Iron chelators, such as Lf, have been repeatedly proposed as a potential therapeutic target during infections [79] and even in COVID-19, we assessed the reduction of ferritin levels during bLf administration, demonstrating its anti-inflammatory activity together with iron chelating ability, which is pivotal for bacterial and viral replication, and at the basis of its antibacterial and antiviral activity[18,20,21].

Liver function is known to be deranged in COVID-19 and a meta-analysis showed that 16% and 20% of patients with COVID-19 had ALT and AST levels higher than the normal range [80]. Liver biochemistry abnormality in COVID-19 patients could be ascribed to several factors, such as direct hepatocyte injury by the virus, drug-induced liver injury, hypoxic-ischemic microcirculation disorder, and underlying liver diseases [69]. In our study, we observed that bLf reduced transaminases levels, decreasing the risk of liver-injury among COVID-19 patients, which is a frequent complication in SARS-CoV-2 severe forms [81].

Regarding clinical symptoms recovery in the bLF arm we observed a gradual reduction of all symptoms, with the exception of fatigue, which persisted in 21.9 % of patients. We explained this result considering patients age and concomitant comorbidities, which could create a bias to identify COVID-19 symptoms. On the other hand, in SOC-treated patients, we observed a partial symptoms recovery and particularly the persistence of anosmia and ageusia in all the affected patients, even at the end of treatment (Figure 8).

Concerning bLf safety, we reported gastrointestinal complaints in 2 patients as occasional findings that did not lead to treatment discontinuation. Therefore, we concluded that bLf is safe and well tolerated among our study population.

Regarding the SOC regimen group we observed limited adverse events related to the assumption of hydroxychloroquine (HCQ), even though its use is controversial as in literature are reported cases of liver injuries linked to the drug [82].

In our analysis, we used formulations containing bLf embedded in liposomes for nasal/oral administration. Indeed, the bLf at 5% of iron saturation form is best suited to obtain the maximum chelating effect. Nucleic digestion, in the nasal cavities, and proteases and lipases hydrolysis, at gastric and intestinal level, inactivate the protein at its first entry, cancelling or extremely reducing the activity. BLf is unstable in water and is particularly sensitive to bacterial and human proteases. This results in protein denaturation, poor absorption and inactivation. The inclusion of bLf in preserving structures, such as liposomes, reduces gastric and intestinal denaturation while maintaining its integrity and therefore its biological functionality[83–85].

One of the limitations of our study was the small sample size of the clinical trial. Further studies, both *in vitro* and *in vivo* are needed to better deepen Lf placement against COVID-19, both as a preventive, adjunctive or in monotherapy. Nevertheless, we achieved a statistical significance in the crucial blood parameters related to disease evolution and we still observed an improving trend in all other analysed markers. Further studies on larger samples are needed to better evaluate the role Lf in treating SARS-CoV-2.

Considering the risk of COVID-19 relapse [86], we also suggest additional long-term studies to evaluate the maintenance of viral clearance with Lf continuous administration.

Lf could be considered as an effective supplement in mild to moderate and asymptomatic COVID19 patients, which are not clearly included in therapeutical guidelines, allowing improvement of patient outcomes and prevention of hospital recovery, but also prevention of the chronic consequences of infection as well as prevention of transmission by shortening the time length of infectiousness.

This study is part of the GEFACOVID2.0 research program coordinated by the Tor Vergata University of Rome.

## Acknowledgements

We thank Prof. Denis Mariano for English language editing.

We thank Technology Dedicated To Care (TDC) for technical support.

The computing resources and the related technical support were provided by CRESCO/ENEAGRID High Performance Computing infrastructure. CRESCO/ENEAGRID High Performance Computing infrastructure is funded by ENEA, the Italian National Agency for New Technologies, Energy and Sustainable Economic Development and by Italian and European research programmes, see http://www.cresco.enea.it/english for information.

