## Supplementary for "Lactoferrin as potential supplementary nutraceutical agent in COVID-19 patients: *in vitro* and *in vivo* preliminary evidences"

### Supplemental data

#### Tables

| A) Spike-Bovine lactoferrin complex simulation. | | | | |
|---|---|---|---|---|
| **VdW (kcal/ mol)** | **Electrostatic (kcal/mol)** | **Nonpolar solvation (kcal/ mol)** | **Polar solvation (kcal/ mol)** | **Interaction energy (kcal/ mol)** |
| -138.27 | 125.11 | -42.29 | -18.35 | -28.02 |
| **B) Spike-Human lactoferrin complex simulation.** | | | | |
| **VdW (kcal/ mol)** | **Electrostatic (kcal/mol)** | **Nonpolar solvation (kcal/ mol)** | **Polar solvation (kcal/ mol)** | **Interaction energy (kcal/ mol)** |
| -158.27 | -115.80 | -21.18 | 209.78 | -48.25 |

**Table S1. A)** Results of the MM/GBSA analyses performed over the last 15 ns of the Spike-Bovine lactoferrin complex simulation**.**

**B)** Results of the MM/GBSA analyses performed over the last 15 ns of the Spike-Human lactoferrin complex simulation**.**

| **Interaction (Spike – Bovine lactoferrin)** | **Interaction type** | **Interaction (Spike Lactoferrin)** | **Interaction type** |
|---|---|---|---|
| Glu159-Arg408 | salt bridge | Lys378-Glu356 | salt bridge |
| Glu162-Lys417 | salt bridge | Asp405-Arg362 | salt bridge |
| Glu355-Arg408 | salt bridge | Asp405-Arg628 | salt bridge |
| | | Glu406-Arg362 | salt bridge |
| Gly404-Glu355 | non-polar contact | Glu406-Arg628 | salt bridge |
| Asp405-Glu355 | non-polar contact | Arg408-Glu355 | salt bridge |
| Arg408-Ala354 | non-polar contact | Arg408-Glu356 | salt bridge |
| Arg408-Glu355 | non-polar contact | Lys444-Glu128 | salt bridge |
| Arg408-Lys358 | non-polar contact | Lys462-Asp449 | salt bridge |
| Arg408-Thr353 | non-polar contact | Glu465-Hie635 | salt bridge |
| Asn437-Gln386 | non-polar contact | Glu465-Lys639 | salt bridge |
| Asn439-Gln386 | non-polar contact | Arg466-Asp646 | salt bridge |
| Gly502-Glu356 | non-polar contact | | |
| Gly502-Thr353 | non-polar contact | Arg403-Arg362 | non-polar contact |
| Val503-Ala359 | non-polar contact | Asp405-Arg362 | non-polar contact |
| Val503-Arg363 | non-polar contact | Arg408-Arg362 | non-polar contact |
| Val503-Glu352 | non-polar contact | Asn437-Asn157 | non-polar contact |
| Val503-Glu355 | non-polar contact | Asn439-Asn127 | non-polar contact |
| Val503-Glu356 | non-polar contact | Asn439-Pro154 | non-polar contact |
| Val503-Thr353 | non-polar contact | Ser443-Asn127 | non-polar contact |
| Gly504-Glu355 | non-polar contact | Ser443-Asn254 | non-polar contact |
| Gly504-Glu356 | non-polar contact | Ser443-Phe155 | non-polar contact |
| Tyr505-Gln386 | non-polar contact | Asn440-Gln130 | non-polar contact |
| Gln506-Gln386 | non-polar contact | Lys444-Asn127 | non-polar contact |
| | | Lys444-Glu128 | non-polar contact |
| Asp405-Ser160 | hydrogen bond | Pro499-Asn254 | non-polar contact |
| Arg434-Glu355 | hydrogen bond | Pro499-Asp253 | non-polar contact |
| Thr526-Ser160 | hydrogen bond | Pro499-Leu126 | non-polar contact |
| Asn527-Ser160 | hydrogen bond | Pro499-Phe155 | non-polar contact |
| Val529-Ser160 | hydrogen bond | Pro499-Pro252 | non-polar contact |
| | | Thr500-Asp253 | non-polar contact |
| | | Thr500-Pro252 | non-polar contact |
| | | Val503-Arg362 | non-polar contact |
| | | Gly504-Arg362 | non-polar contact |
| | | Tyr505-Pro154 | non-polar contact |
| | | Tyr508-Thr159 | non-polar contact |
| | | Asp405-Arg362 | hydrogen bond |
| | | Arg408-Glu356 | hydrogen bond |
| | | Asn437-Asn157 | hydrogen bond |
| | | Asn437-Pro154 | hydrogen bond |
| | | Asn439-Pro154 | hydrogen bond |
| | | Asn440-Gln130 | hydrogen bond |
| | | Ser443-Asn127 | hydrogen bond |
| | | Lys444-Asn127 | hydrogen bond |

| | | |
|---|---|---|
| | Lys444-Glu128 | hydrogen bond |
| | Thr500-Pro252 | hydrogen bond |

**Table S2 (LEFT SIDE)** Molecular interactions established between the RBD domain of the Spike protein and the Bovine lactoferrin. Only interactions identified for more than 50% of simulation time are shown. Residues highlighted in grey are shared in the interfaces of Spike-ACE2 and Spike-Bovine lactoferrin.

**(RIGHT SIDE)** Molecular interactions established between the RBD domain of the Spike protein and the lactoferrin. Only interactions identified for more than 50% of simulation time have been showed. Residues highlighted in grey are shared in the interfaces of Spike-ACE2 and Spike-Human lactoferrin

| A) Statistical analysis evaluation between COVID-19 bLf group and control group | | | | |
|---|---|---|---|---|
| | **COVID-19 bLf patient platelet count (10^3/ ul)** | **Healty control platelet count (10^3/ ul)** | **COVID-19 bLf patient neutrophilis count (10^3/ ul)** | **Healty control neutrophilis count (10^3/ ul)** |
| **Mean** | 239,63 | 255,97 | 4,46 | 5,84 |
| **Variance** | 6897,09 | 4652,45 | 5,66 | 113,06 |
| **standard deviation** | 179324,34 | 134921,05 | 113,2 | 3278,74 |
| **t _Student** | | -3,07 | | -2,05 |
| **two-tailed p-value** | | < 0,0001 | | 0,04 |
| | **COVID-19 bLf patient monocytes count (10^3/ ul)** | **Healty control monocytes count (10^3/ ul)** | **COVID-19 bLf patient D-Dimer (ng/ml)** | **Healty control Ddimer (ng/ml)** |
| **Mean** | 0,5 | 0,69 | 1546,82 | 484 |
| **Variance** | 0,04 | 1,22 | 13034194,25 | 420241,48 |
| **standard deviation** | 0,8 | 35,38 | 273718079,3 | 11346519,96 |
| **t _Student** | | -2,73 | | 5,37 |
| **two-tailed p-value** | | 0,006 | | < 0,0001 |
| | **COVID-19 bLf patient AST (U/L)** | **Healty control AST (U/L)** | **COVID-19 bLf patient Ferritin (ng/ml)** | **Healty control Ferritin (ng/ml)** |

| | | | | |
|---|---|---|---|---|
| **Mean** | 24,36 | 26,17 | 201,43 | 119,24 |
| **Variance** | 95,99 | 76,35 | 57729,26 | 23855,76 |
| **standard deviation** | 2303,76 | 2214,15 | 1039126,68 | 429403,68 |
| **t _Student** | | -2,67 | | 3,87 |
| **two-tailed p-value** | | 0.008 | | < 0,0001 |
| | **COVID-19 bLf patient lymphocytes count (10^3/ ul)** | **Healty control lymphocytes count (10^3/ ul)** | **COVID-19 bLf patient ALT (U/L)** | **Healty control ALT (U/L)** |
| **Mean** | 1,75 | 3,06 | 29,36 | 26,8 |
| **Variance** | 0,3 | 27,38 | 515,49 | 229,13 |
| **standard deviation** | 6 | 794,02 | 12371,76 | 6644,77 |
| **t _Student** | | -4 | | 1,84 |
| **two-tailed p-value** | | < 0,0001 | | 0,059 |
| | **COVID-19 bLf patient Adrenomedullin (pg/ml)** | **Healty control Adrenomedullin (pg/ml)** | **COVID-19 bLf patient IL-6 (pg/ml)** | **Healty control IL-6 (pg/ml)** |
| **Mean** | 0,57 | 0,42 | 38,1 | 7,48 |
| **Variance** | 0,19 | 0,01 | 13232,16 | 0,03 |
| **standard deviation** | 1,71 | 0,15 | 198482,4 | 0,45 |
| **t _Student** | | 3,32 | | 3,01 |
| **two-tailed p-value** | | < 0,0001 | | < 0,0001 |

**B) Statistical analysis evaluation between T2 and T0 in COVID-19 bLf group**

| Statistical analysis | **D-Dimer (ng/ml) $\Delta(T_2 - T_0)$** | **Ferritin (ng/ml) $\Delta(T_2 - T_0)$** | **IL-6 (pg/ml) $\Delta(T_2 - T_0)$** |
|---|---|---|---|
| Mean | -392,56 | -90,63 | -2,52 |
| Variance | 366600,7 | 42319,88 | 36,31 |
| Standard Deviation | 142,71 | 48,49 | 1,46 |
| t-Student | -2,75 | -1,87 | -1,73 |

| | | | |
|---|---|---|---|
| p-value | 0,01 | 0,04 | 0,05 |

| **C) Statistical analysis evaluation between T2 and T0 in COVID-19 SOC group** | | | |
|---|---|---|---|
| Statistical analysis | **Hemoglobin (g/dl) $\Delta(T_2 - T_0)$** | **Red Blood Cells count (10^6/ ul) $\Delta(T_2 - T_0)$** | **Platelets count (10^3/ ul) $\Delta(T_2 - T_0)$** |
| Mean | -2,22 | -0,43 | 89,56 |
| Variance | 14,57 | 0,46 | 5713,33 |
| Standard Deviation | 14,76 | 2,63 | 292,75 |
| t-Student | -2,337 | -2,529 | 4,739 |
| p-value | 0,05 | 0,05 | 0,001 |

**Tab. S3 A)** Statistical analysis evaluation between COVID-19 bLf group and control group**; B)** Statistical analysis evaluation between T2 and T0 in COVID-19 bLf group**; C)** Statistical analysis evaluation between T2 and T0 in COVID-19 SOC group

**Figures**

CONSORT
TRANSPARENT REPORTING of TRIALS

### CONSORT 2010 Flow Diagram

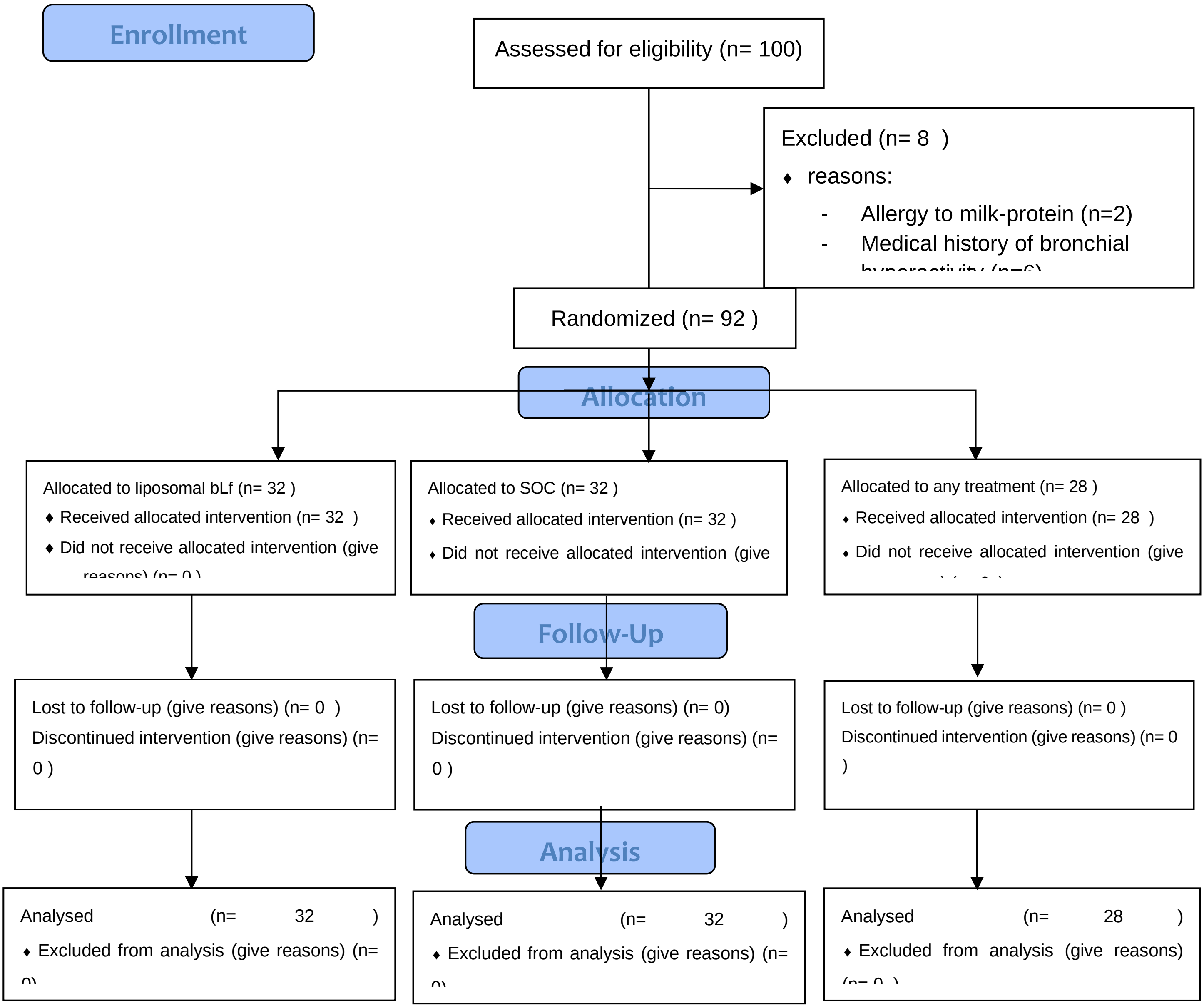

**Fig. S1:** CONSORT diagram of the clinical trial

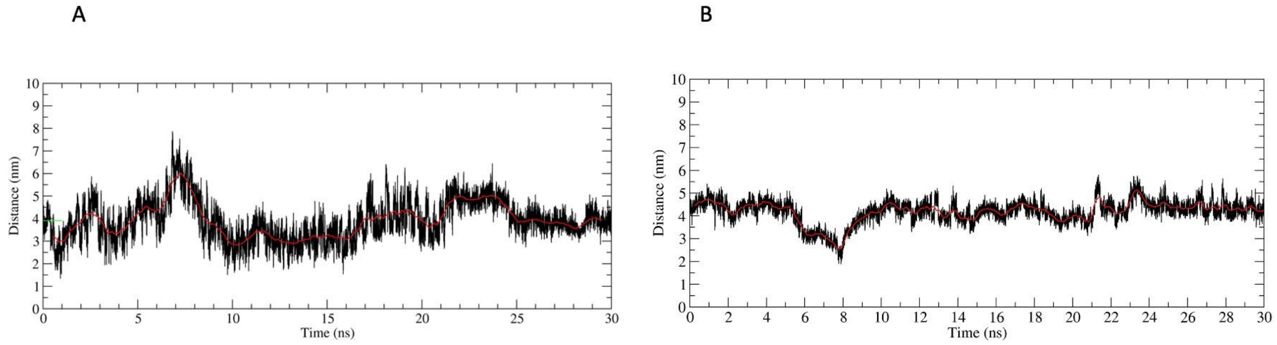

**Figure S2. A)** Time-dependent analysis of the distance evaluated between the centers of mass of the CTD1 domain in the up conformation and of the bovine lactoferrin. The red line represents the distance values averaged over 250 trajectory frames.

**B)** Time-dependent analysis of the distance evaluated between the centers of mass of the CTD1 domain in the up conformation and of the human lactoferrin. The red line represents the distance values averaged over 250 trajectory frames.

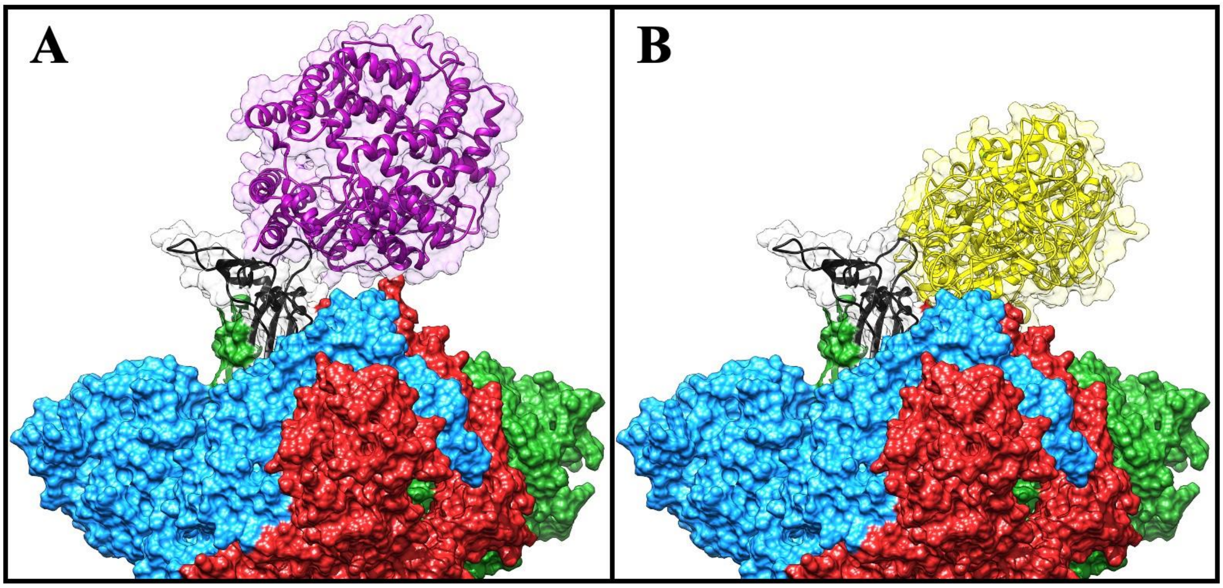

**Figure S3.** Structural superposition of the Frodock best complex and of the ACE2-Spike glycoprotein (PDB ID: 6LZG). The red, blue and green solid surfaces represent the three different chains composing the Spike glycoprotein. The black ribbons highlight the CTD1 domain in the up conformation. The magenta and yellow ribbons represent the ACE2 (A) and the human lactoferrin (B), respectively, surrounded by a transparent molecular surface representation, in order to point out the positions occupied in the space by the different structures
